# Towards customized studies of G-quadruplex DNA structures in live cells

**DOI:** 10.1101/2021.04.07.438780

**Authors:** Bagineni Prasad, Mara Doimo, Måns Andréasson, Valentin L’Hôte, Erik Chorell, Sjoerd Wanrooij

**Author notes:** Electronic Supplementary Information (ESI) available: [Experimental procedures, compound evaluations, and characterization of the compounds]. See DOI: 10.1039/x0xx00000x. These authors contributed equally to this work (Bagineni Prasad, Mara Doimo).

## Abstract

G-quadruplex (G4) DNA structures are implicated in central biological processes and are considered promising therapeutic targets because of their links to human diseases such as cancer. However, functional details of how, when, and why G4 DNA structures form *in vivo* are largely missing leaving a knowledge gap that requires tailored chemical biology studies in relevant live-cell model systems. Towards this end, we developed a synthetic platform centered around one of the most effective and selective G4 stabilizing compounds, Phen-DC3. We used a structure-based design to equip Phen-DC3 with an amine in a position that does not interfere with G4 interactions. To evaluate the power of the approach, we next used this reactive handle to conjugate a BODIPY fluorophore and a cell-penetrating peptide to Phen-DC3. The BODIPY conjugation generated a fluorescent derivative with retained G4 selectivity, G4 stabilization, and cellular effects that revealed the localization and function of Phen-DC3 in human cells. On the other hand, the cell-penetrating peptide conjugation, while retaining G4 selectivity and stabilization, increased nuclear localization and cellular effects, showcasing the potential of this approach to modulate and direct cellular uptake e.g. as delivery vehicles. The developed platform can thus generate tailored biochemical tools for the studies of G4 biology to uncover molecular details and therapeutic approaches.

## Introduction

G-quadruplex (G4) DNA structures are four-stranded DNA structures that lately have gained significant scientific interest because of their links to key cellular events. The first report related to G4 DNA structures dates back to 1910 when Ivar Christian Bang mentioned that guanylic acid can form a gel at higher concentrations.^1^ In 1962, Martin Gellert *et al* followed up on this work and published the first structure of a G-tetrad.^2^ At that time, the G4 DNA structure was believed to be an *in vitro* artefact and research related to G4 DNA was for long not prioritized. However, a growing body of evidence has lately shown that G4 DNA structures are highly abundant, evolutionary conserved, and seem to play important roles in the processing of our genetic information.^3, 4^ The core of a G4 structure consists of layers of guanines that stack on each other. Each guanine layer, called a G-quartet or G-tetrad, consists of four guanines that bind to each other by Hoogsteen hydrogen bonding.^2, 5^ The G4 structures are further stabilized by monovalent cations,^6–8^ and can altogether form a structure that is more stable than double stranded DNA (dsDNA).^9^ However, the stability of a G4 structure is highly dependent on the direction of the four G-strands that form the central guanine core and the composition, length, and position of the nucleotide sequences connecting the G-strands, called loops. Although the guanine core structure is similar between G4 structures, the loops can be highly variable and thus confer large structural diversity between G4 structures.^10–12^ Furthermore, G4 structures can be formed intramolecular (by one DNA strand) or intermolecular (by two or four individual strands) and present a diversity of topologies as defined by the parallel or antiparallel strand orientation.^13^ Bioinformatics studies have revealed more than 700,000 motifs with the potential to form G4 structures.^14^ Their location is distributed throughout the human genome, but they are enriched at promoters and specifically in the promoters of many oncogenes such as *c-MYC, BCL2, VEGF*, and *c-KIT*.^15, 16^ G4 structures are thus considered promising drug targets and are investigated in different therapeutic strategies to treat various cancers.^17–19^ The abundance of G4 structures and their enrichment at certain genomic locations are strong indications of their cellular importance. Studies have identified potential regulatory roles of G4s in replication, transcription, splicing, translation, and recombination.^4, 14^ In addition to this, G4 structures also impact the mitochondrial DNA (mtDNA) gene expression. Putative G4-forming sequences are found all around the 16kb-long circular DNA molecule that shapes the human mitochondrial genome,^20, 21^ moreover G4 structures have a functional role during mitochondrial transcription.^22, 23^ Nonetheless, the precise influence of G4 structures on mtDNA maintenance is still largely unknown. Taken together, there are vast amounts of G4 structures that possess broad structural diversity, and their presence is linked to key biological processes. However, there are still considerable gaps in the knowledge of G4 biology and new techniques and strategies to study G4 biology are of immense interest.

Chemical biology is a multidisciplinary research field at the interface of chemistry and biology that generally involves manipulation and studies of biology using small molecules. One subcategory of chemical biology concerns the generation and validation of new small-molecule-based tool compounds. These serve as excellent tools to modulate biological systems and possess unique characteristics that have proven highly valuable to e.g. study G4 biology. Indeed, one way to study G4 DNA structures is to develop small molecules that bind and stabilize G4 structures. Phen-DC3 is one of the most frequently used and strongest G4 stabilizing compounds.^24^ However, to allow for tailored studies of G4 structures one would require the possibility to functionalize Phen-DC3 with different handles for customized studies such as pull-downs, fluorescence studies, target delivery peptides, etcetera. Here, we have developed a synthetic platform based on Phen-DC3 that allows for such customized functionalization. The utility of this synthetic platform is exemplified by the introduction of a 4,4-Difluoro-4-bora-3a,4a-diaza-*s*-indacene (BODIPY) fluorophore that revealed the basis for the limited cellular effect of Phen-DC3 despite its strong G4 binding and stabilization. Next, the diversity of the approach is further exemplified by the conjugation of a cell-penetrating peptide that effectively opens the possibility for cellular studies. Importantly, even though the conjugated functionalities are large and charged, their presence does not alter the potent and selective G4 stabilization of Phen-DC3. These studies thus show the power of the presented strategy to enable further chemical biology studies aimed at G4 biology.

## Results and discussions

Phen-DC3 is one of the strongest G4 stabilizers reported (Figure 1A). *In vitro*, Phen-DC3 can stabilize with high affinity and specificity G4 structures of different topologies.^24, 25^ However, in HeLa cells, the effect of Phen-DC3 on cell viability is surprisingly mediocre, with only 20% cell death after 100 μM treatment for 48h (Figure 1B).

**Figure 1.**
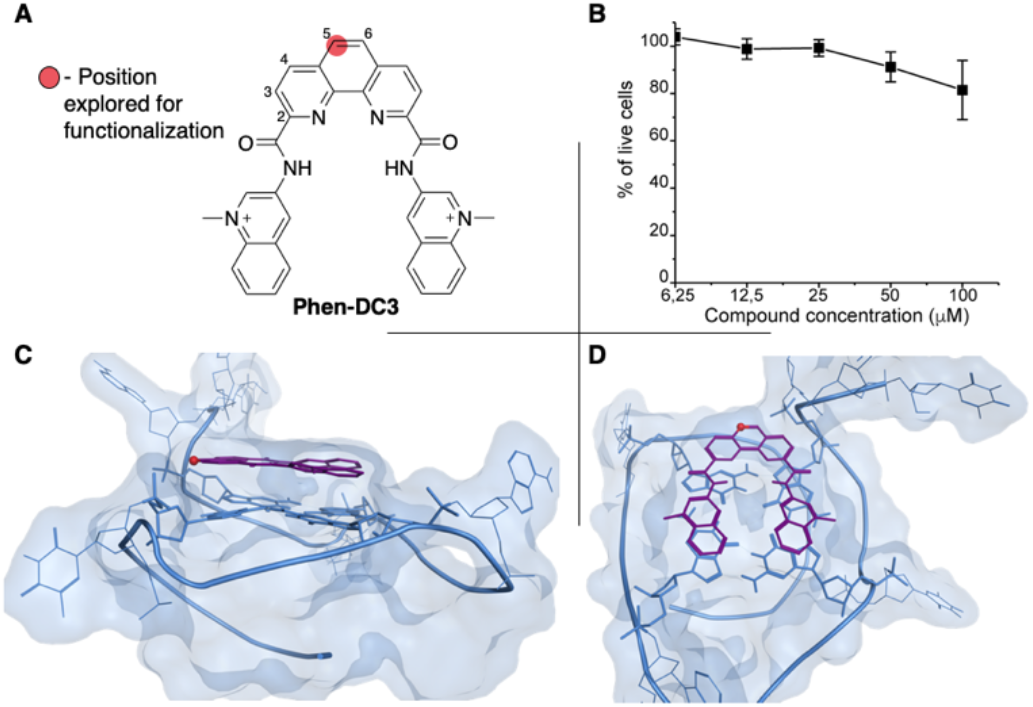
**A** Chemical structure of Phen-DC3. **B** Resazurin-based cell viability assay on Hela cells treated for 48h with Phen-DC3 at the indicated concentrations. Data represent mean ± standard deviation of three independent experiments. **C** and **D** Crystal structure (2MGN) of Phen-DC3 (purple) and Pu24T (blue) viewed from the side (**C**) and top (**D**). The carbon labelled with a red sphere in **A**, **C**, and **D** represents the position on the phenanthroline core suitable for functionalization.

We hypothesized that this is correlated to low cellular uptake that hampers the utility of this commercially available, well-characterized, and recognized G4 stabilizing compound in cellular studies of G4 biology. To address this, we set out to develop a synthetic platform that would allow for the functionalization of Phen-DC3 with various versatile substituents for customized cellular studies without affecting the integrity of Phen-DC3 and its very strong and selective G4 stabilization. For this purpose, we investigated reported structural data on how Phen-DC3 interacts with G4 DNA structures, *e.g*. the structure of Phen-DC3 bound to the Pu24T *c-MYC* G4 DNA structure.^26^ This revealed positions 5 and 6 on the central phenanthroline scaffold as excellent connection points for further functionalization of Phen-DC3 without affecting the G4 binding properties (Figure 1C and 1D). Hence, the preparation of a Phen-DC3 derivative with a primary amine in this position would open up for further functionalization. To achieve this, a selective nitration of Neocuproine **1** using a combination of fuming sulfuric acid and nitric acid at 120 °C was used to generate nitro derivative **2** (Scheme 1). Compound **2** was next converted to dicarbaldehyde **3** by selenium mediated oxidation, which was further oxidized to the dicarboxylic acid derivative **4** in 56% overall yield.^27^ A coupling reaction between acid **4** and **5** (3-aminoquinoline) generated compound **6** in 87% yield using 1-[bis(dimethylamino)methylene]-1H-1,2,3-triazolo[4,5-b]pyridinium 3-oxide hexafluorophosphate (HATU) and *N*,*N*-Diisopropylethylamine (DIPEA) at room temperature. Selective reduction of the nitro functionality in compound **6** without affecting the quinoline group was subsequently performed using a Pd/C mediated hydrogenation to give the desired primary amine **7** in 72% yield.

**Scheme 1.**
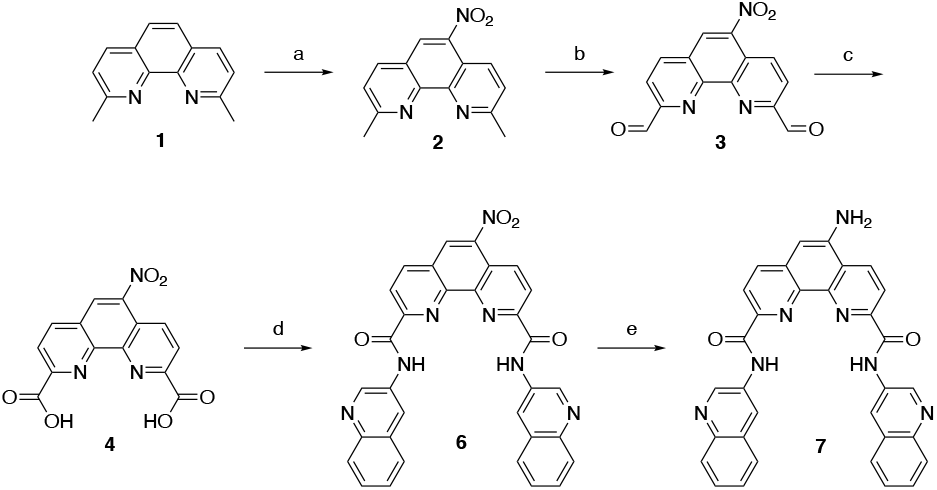
Synthesis of amino substituted Phen-DC3. Conditions: a) HNO_3_, H_2_SO_4_, 120 °C, 1 h, 44%; b) SeO_2_, 1,4-dioxane:H_2_O (0.4%), reflux, 3 h; c) Conc. HNO_3_ reflux 6 h, (2 steps 56%); d) 3-aminoquinoline **5**, HATU, DIPEA, DMF, rt, 12 h, 87%; e) Pd/C, H_2_, 80 °C, DMF, 3 h, 72%;

To investigate the cellular localization of Phen-DC3 in live cells, we used the developed synthetic scheme to link a BODIPY fluorophore to Phen-DC3 (Scheme 2). The BODIPY fluorophore was chosen because of its high quantum yield of fluorescence, absorption/emission wavelengths, photostability, and relatively small size. Hence, bromoacetyl chloride was reacted with amine **7** in the presence of DIPEA to give bromoderivative **8** which upon treatment with sodium azide provided compound **9**. Staudinger reduction conditions reduced the azido derivative **9** to amine derivative **10** and a subsequent HATU mediated coupling reaction with BODIPY acid^28^ **11** successfully provided the BODIPY linked Phen-DC3 **13** after methylation using methyl iodide.

A Taq polymerase stop assay was next performed to investigate if the addition of the BODIPY fluorophore would affect the ability of Phen-DC3 to selectively stabilize G4 DNA. This assay measures how efficiently the Taq polymerase can synthesize new DNA from a template strand with a single nucleotide resolution readout by extending a fluorescent 5’ end-labelled DNA primer. A G4 DNA structure is an obstacle for the DNA polymerase, and if a template strand with a G4 DNA structure is used, the Taq polymerase will halt one or two nucleotides before the first G-tetrad in the G4 structure.

**Scheme 2.**
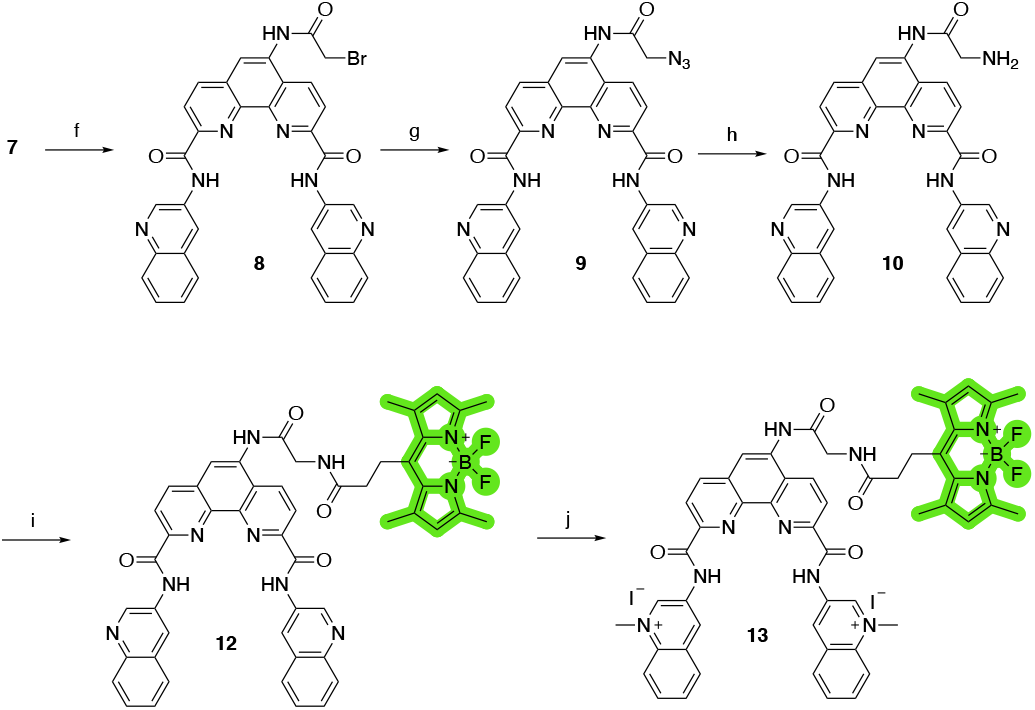
Synthesis of Phen-DC3 linker with BODIPY. Conditions: f) Bromoacetyl chloride, Et_3_N, rt, 4 h, 80%; g) NaN_3_, DMF, rt, 70%; h) PPh_3_, THF, H_2_O, 90 °C, 8 h, 72%; i) 3-(5,5-difluoro-1,3,7,9-tetramethyl-5H-4λ4,5λ4 di-pyrrolo[1,2-c:2’,1’-f][1,3,2]diaza-borinin-10-yl)propanoic acid **11**, HATU, DIPEA, DMF, rt, 1 h, 65%; j) MeI, DMF, 40 °C, 24 h, 70%.

However, the DNA polymerase can fairly easily resolve the G4 structure and synthesize the full-length run-off DNA product. The efficiency of G4 stabilizing compounds can be measured by their ability to increase the stalling of the DNA polymerase at the G4 structure on the template strand. Also, it provides nucleotide resolution of the DNA polymerase stalling. We performed these primer extension assays in the presence of DMSO (neg. control), Phen-DC3 alone, BODIPY alone, and Phen-DC3 linked to BODIPY using a G4 (Figure 2A) and a non-G4 (Figure 2B) containing DNA template. In the absence of a G4 stabilizer (DMSO) the DNA polymerase could extend the DNA primer beyond the G4 structure in the DNA template to generate a full-length run-off DNA product (Figure 2A, lane2). As expected, the addition of Phen-DC3 led to a strong stabilization of the template G4, resulting in a clear DNA polymerase stalling before the G4 structure (Figure 2A lane 3). Interestingly, the BODIPY linked Phen-DC3 retained its G4 stabilization ability (Figure 2A, lane 4), whereas the BODIPY fluorophore alone did not affect G4 formation (Figure 2A lane 5).

**Figure 2:**
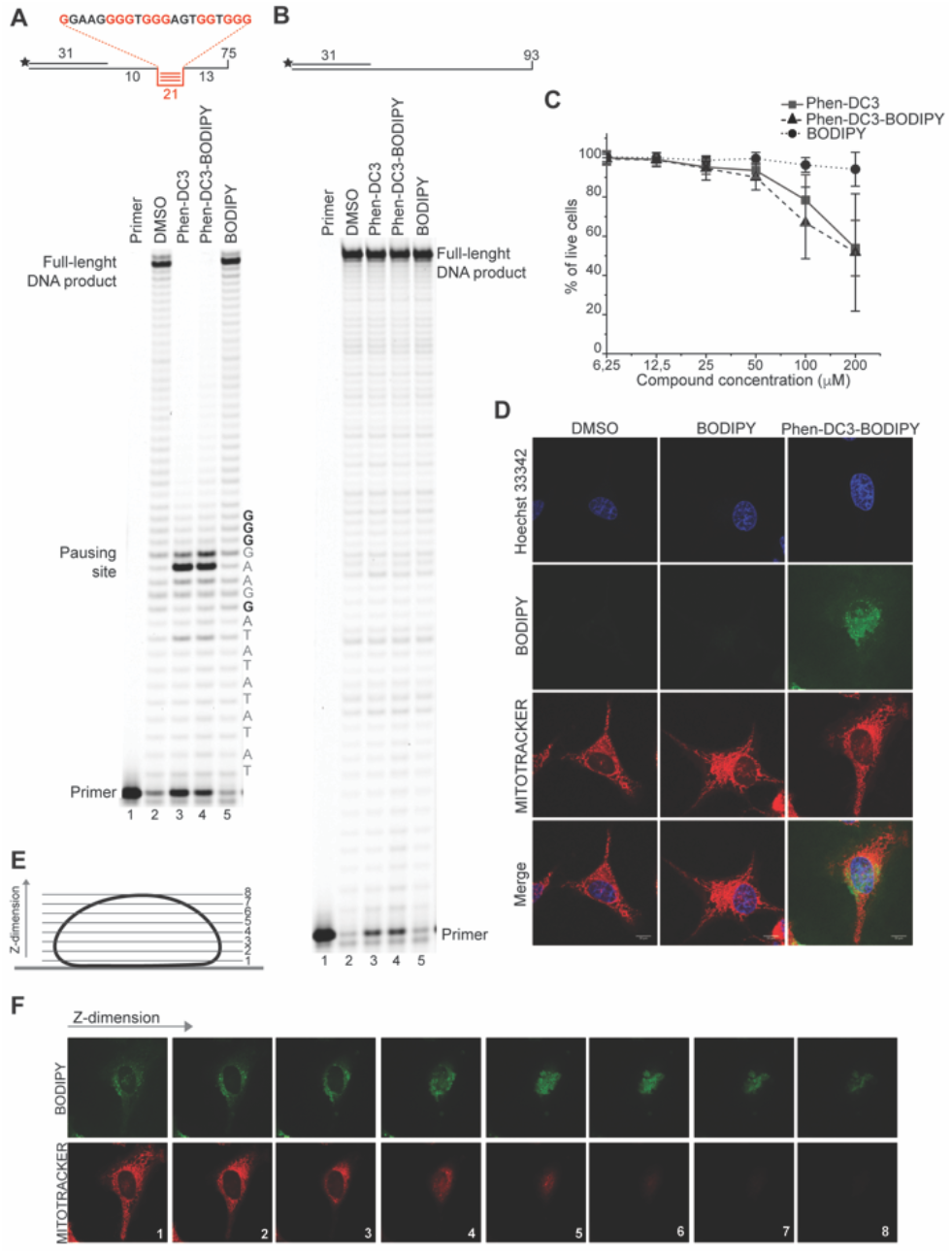
Phen-DC3-BODIPY has the same effect than Phen-DC3 *in vitro* and in HeLa cells and does not enter the nucleus. **A.** Taq-polymerase STOP assay on a template with a G4 structures (as depicted above the gel) in the presence of 1μM of the indicated compounds. The nucleotides of the template upstream and around the pausing site are indicated. Guanines involved in the formation of the G4 are indicated in bold. **B.** Taq-polymerase STOP assay on a non-G4 DNA template in the presence of 1μM of the indicated compounds. **C.** Resazurin-based cell viability assay on Hela cells treated for 48h with Phen-DC3, Phen-DC3-BODIPY or BODIPY at the indicated concentrations in high glucose medium. Data represent mean ± standard deviation of three independent experiments. **D.** Maximum intensity projections of confocal images of HeLa cells treated for 12h with 5uM of the indicated compounds. Cells were trypsinized, washed and allowed to seed again in on new coverslip for approximately 4h before imaging. Mitochondria were stained with Mitotracker red, nuclei were stained with Hoechst 33342. **E.** Schematic representation of the Z–stacks acquisition at the confocal microscopy. **F**. Single Z-stacks of HeLa cells treated with 5μM Phen-DC3-BODIPY. Mitochondria were stained with Mitotracker red. The numbers indicate the single stacks as represented in scheme E.

Similar to Phen-DC3 alone, the BODIPY linked Phen-DC3 retained its G4 selectivity and displayed no effect in the DNA polymerase stop assay when a template strand without G4 structure was used in the reaction (Figure 2B, lane 3 and 4).

To further verify that the BODIPY linking strategy does not affect the cellular properties of Phen-DC3, we performed cell viability studies, which revealed near to identical effects of Phen-DC3-BODIPY compared to treatment with Phen-DC3 alone (Figure 2C). BODIPY alone had minimal effects on cell viability (Figure 2C). We conclude that introducing the BODIPY fluorophore to Phen-DC3 using our synthetic platform has generated a fluorescent Phen-DC3 derivative that has retained *in vitro* and cell viability effects of Phen-DC3. This allowed us to use fluorescence live-cell imaging experiments to study the localization of this compound. Cells were treated with 5 μM of BODIPY linked Phen-DC3 for 12 h, washed, and allowed to adhere for a period of 6 h before live-cell studies (Figure 2D).

This revealed that Phen-DC3-BODIPY does enter the cells but, strikingly, that there is no presence of BODIPY linked Phen-DC3 in the nucleus (no-colocalization with Hoechst 33342 (nuclear) staining). This fact could explain the minimal effect of Phen-DC3 on cell viability, despite it being a highly selective and efficient G4 stabilizer that should have detrimental effects when localized to the nucleus (Figure 1B and 2C). Interestingly, some overlap with the mitochondrial staining (Mitotracker) is observable, suggesting that Phen-DC3 might enter mitochondria (Figure 2E and F). Based on this observation, we hypothesized that a mitochondrial localization of Phen-DC3 should lead cells to be more sensitive when maintained in growth conditions that depend on mitochondrial function. Indeed, HeLa cells were more sensitive to Phen-DC3 treatment when dependent on mitochondrial energy production (galactose or low glucose medium) compared to highly glycolytic cells (grown in high glucose-medium) (Compare Figure 1B to Figure 3A and 3B). The Phen-DC3-BODIPY derivative showed identical results compared to Phen-DC3 alone, which again underlines the power of the developed synthetic platform.

**Figure 3:**
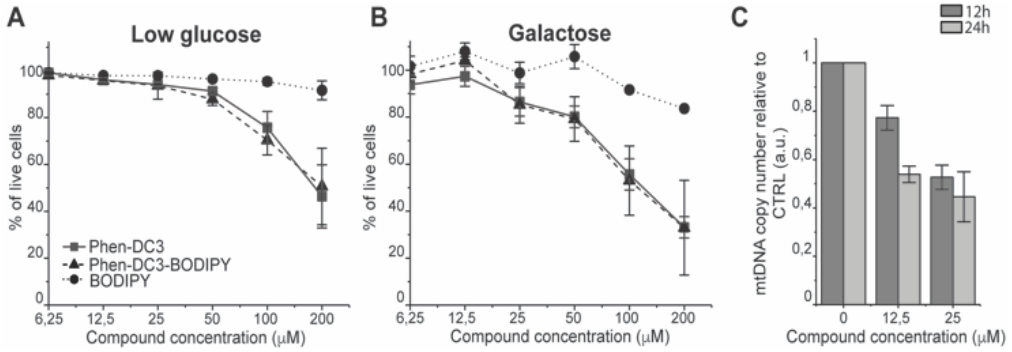
Phen-DC3 affects mtDNA copy number in HeLa cells. Resazurin-based cell viability assay on Hela cells treated for 48h with Phen-DC3, Phen-DC3-BODIPY or BODIPY at the indicated concentrations in low glucose medium (A) or in galactose medium (**B**). Data represent mean ± standard deviation of three independent experiments. **C.** mtDNA copy number of HeLa cells treated for 12h or 24h with Phen-DC3 at the indicated concentrations. MtDNA copy number was determined by quantitative PCR from total DNA. The relative mtDNA copy number was expressed as the ratio between mtDNA ATP6 region and nuclear DNA r18S region. Data represent mean ± abs error of two independent experiments.

Because Phen-DC3 is highly selective for G4 DNA binding and stabilization *in vitro*, we hypothesize that its target in the mitochondria is the mtDNA and its potential G4 structures. Formation and stabilization of G4 structures can lead to stalling of the mtDNA replication machinery and partial loss of the total mtDNA pool. To investigate this, we looked at the effect of Phen-DC3 treatment on the mtDNA copy number in HeLa cells. We detected a clear Phen-DC3 concentration-dependent decrease in the mtDNA copy number after 12 h treatment (Figure 3C). Extended Phen-DC3 treatment (24 hours) resulted in a further decreased mtDNA copy number (Figure 3C). We conclude that Phen-DC3 can enter the mitochondria and bind to G-quadruplex structures on the mtDNA to impede mtDNA replication and reduce the mtDNA copy number. Taken together, our data suggest that Phen-DC3 has limited cellular uptake and is unable to enter the nucleus in live cells. The small amount of Phen-DC3 that is taken-up remains cytosolic and exerts its effect on the mitochondrial function by affecting the mtDNA maintenance process.

To increase the ability of Phen-DC3 to enter human cells and thus greatly expand its value as a research tool, we again turned to the developed synthetic platform with the aim to conjugate a cell-penetrating peptide to Phen-DC3, Phen-DC3-PP, with the goal to increase cellular uptake of the compound. The central amine **7** was thus coupled with methyl adipoyl chloride **14** in the presence of 1,5,7-Triazabicyclo[4.4.0]dec-5-ene (TBD) base to give methyl-6-oxohexanoate compound **15** in 82% yield. Ester hydrolysis to hexanoic acid derivative **16** was performed with lithium hydroxide in DMF in 76% yield followed by methylation using methyl iodide in DMF at 40 °C to give the target compound **17** in 92% yield (Scheme 3).

**Scheme 3.**
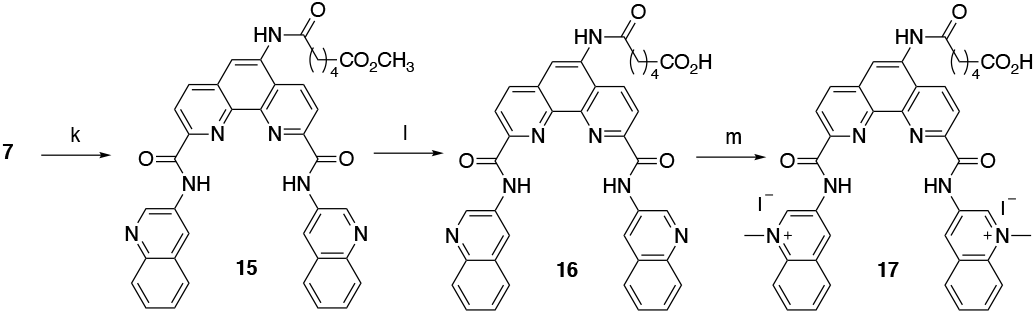
Synthesis of Phen-DC3 linker with acid. Conditions: k) methyl 6-chloro-6-oxohexanoate **14**, TBD, DCM, rt, 3 h, 82%; l) 1M LiOH, DMF, 3 h, 76%; m) MeI, DMF, 40 °C, 18 h, 92%.

The key Phen-DC3 derivative **17** was next linked with the PP as outlined in Scheme 4. The couplings were performed on rink amide resin with 2,2,4,6,7-pentamethyldihydrobenzofuran-5-sulfonyl (Pbf) protected arginine (**18**) using amino acids cyclohexyl alanine **19** and arginine **20**. HBTU and HOBT were used as coupling agents and DIPEA as the base in DMF on a solid phase synthesizer. Acid derivative **17** was subsequently coupled to the *N*-terminus of the resin-bound PP^29^ (rF_x_rF_x_rF_x_r; where r = d-arginine and F_x_ = L-cyclohexylalanine) with HATU as the coupling reagent and DIPEA as the base in DMF. The target Phen-DC3 linked to PP (**22**) was cleaved from the resin with a TFA:TIPS:H_2_O (15:1:1) mixture and then precipitated by the addition of cold diethyl ether (Scheme 4).

**Scheme 4.**
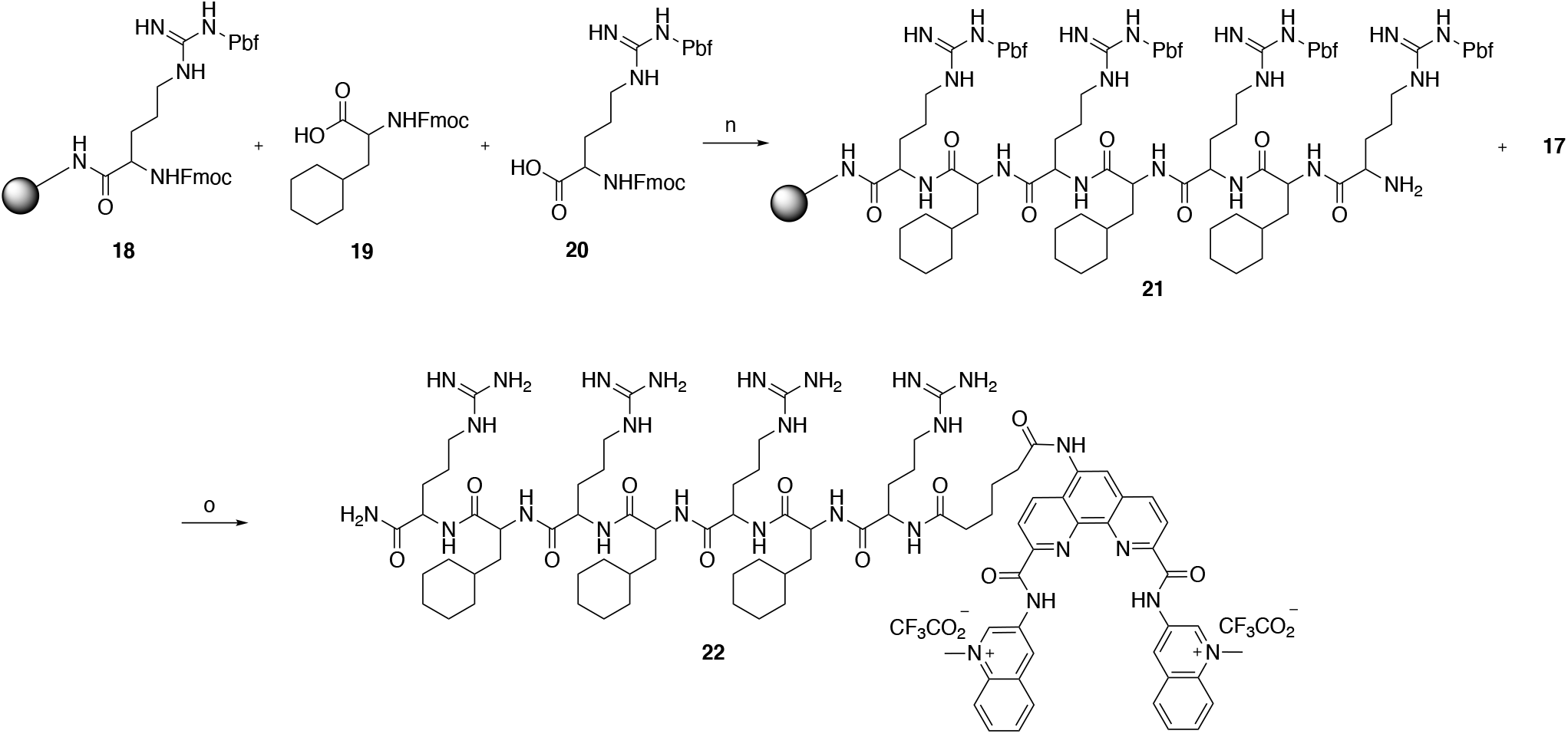
Synthesis of Phen-DC3 linker with PP. Conditions: n) (i) HBTU, HOBT, DIPEA, DMF, rt; (ii) Piperidine; o) (i) **17**, HATU, DIPEA, DMF, rt, 12 h, 87%; (ii) TFA:TIPS:H_2_O (15:1:1), rt, 2h.

To assess if the charged PP addition would affect the ability of Phen-DC3 to selectively stabilize G4 DNA, we performed a Taq DNA polymerase stop assay. Phen-DC3-PP (**22**) retains the ability to stabilize G4 DNA and blocks the DNA polymerase just upstream of the first G-tetrad on the G4 containing DNA template (Figure 4A lanes 26-32). Compared to Phen-DC3 alone and Phen-DC3 with only the linker (supporting Scheme S1), the attachment of the large and charged PP does have a surprisingly low effect on G4 DNA stabilization, and only a slight difference in the G4 stabilization is observed (Figure 4A compare lanes 10-24 with lanes 26-32 and Supporting information figure 1A). Although in the presence of Phen-DC3 some DNA products were detected that had extended DNA beyond the G4, this full-length run-off DNA product signal was strongly reduced compared to the negative control DMSO (6% vs 38% after 30’) (Figure 4B). Importantly, Phen-DC3-PP also retains its selectivity towards G4 and does not affect DNA polymerase extension when a non-G4 DNA template is used (Supporting information figure 1B and 1C).

**Figure 4:**
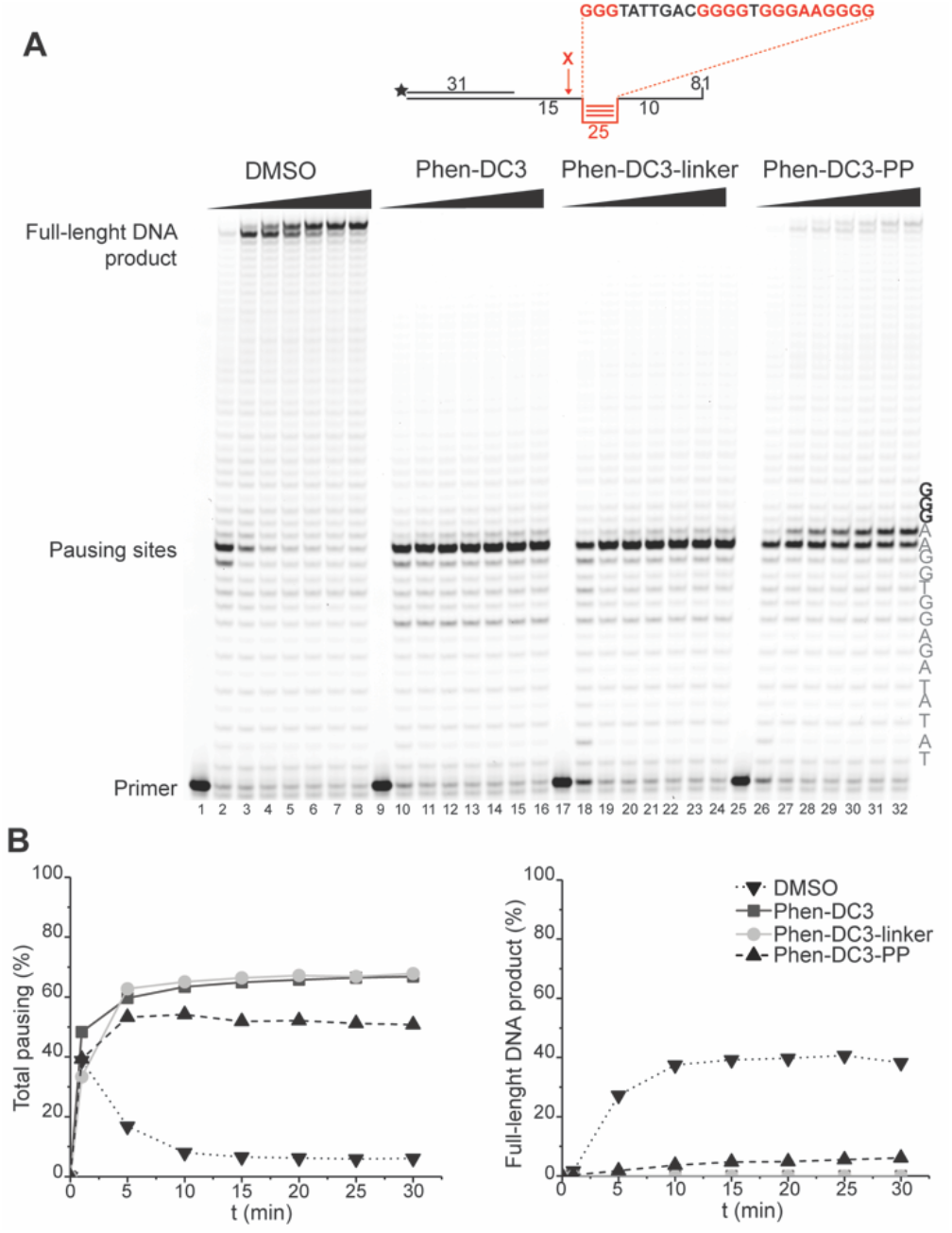
Fusion of the penetrating peptide increases toxicity in HeLa cells. **A.** Taq-polymerase STOP assay on a template with a G4 structure in the presence of 250nM of the indicated compounds. For each compound, the reaction was blocked at increasing time points. The nucleotides of the template upstream and around the pausing site are indicated. Guanines involved in the formation of the G4 are indicated in bold. **B.** Quantification of pausing sites (left) and full-length product (right). Values are represented as % of the total line signal.

To investigate how the ability to stabilize G4 DNA in vitro translates into their effect on cells, we next performed cell viability assays. Indeed, Phen-DC3-PP has a significantly increased effect on cell viability compared to Phen-DC3 alone, which is in line with an increased cellular uptake of PP-linked Phen-DC3 (Figure 5A).

**Figure 5:**
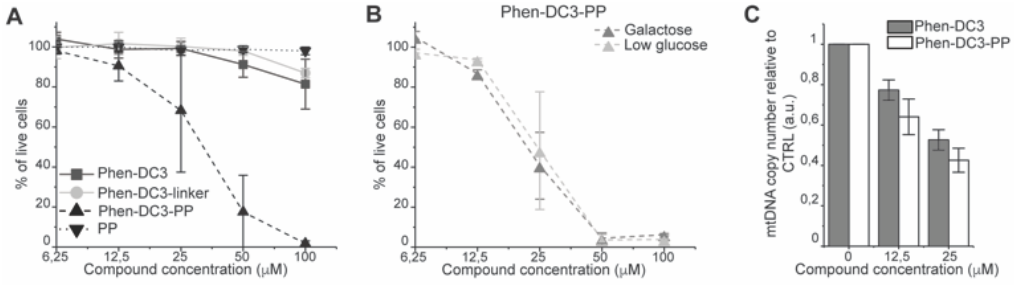
Addition of the penetrating peptide cause DNA damage in HeLa cells. Cell viability assay on HeLa cells treated for 48h with the compounds at the indicated concentrations in high glucose medium (**A**) or in low glucose or galactose media (**B**). Data represent mean ± standard deviation of three independent experiments. **C.** mtDNA copy number of HeLa cells treated for 12h with Phen-DC3 or Phen-DC3-PP at the indicated concentrations. MtDNA copy number was determined by quantitative PCR from total DNA. The relative mtDNA copy number was expressed as the ratio between mtDNA ATP6 region and nuclear DNA r18S region. Data represent mean ± abs error of two independent experiments.

Importantly, the observed effects on cell viability are linked to Phen-DC3 and not to the cell-penetrating peptide as the peptide alone did not have any effect on cell viability (Figure 5A). The stronger cytotoxicity of PhenDC3-PP did not correlate with an increased effect on mitochondrial function as cells treated with Phen-DC3-PP behaved similarly to those treated with Phen-DC3 alone when grown in conditions in which mitochondrial energy production is essential (low glucose or galactose-medium) (Figure 5B). Furthermore, the effect on mtDNA copy number was also comparable between Phen-DC3-PP and Phen-DC3 alone treated HeLa cells (Figure 5C). The increased effect on cell viability by conjugation of a cell-penetrating peptide can therefore possibly be explained by the ability of Phen-DC3-PP to enter the cell nucleus and stabilize nuclear G4 DNA. Conjugation of a cell-penetrating peptide to Phen-DC3 thus increases the cell penetration and the cytotoxic effect of Phen-DC3. This strategy enables studies of Phen-DC3 and its effect on nuclear DNA in cells, which is of high relevance considering that Phen-DC3 is among the most efficient and frequently used G4 DNA stabilizing compounds.

## Conclusions

Here, we have developed a synthetic platform that allows for the functionalization of the well-characterized G4-stabilizer Phen-DC3. We first linked a fluorophore to Phen-DC3 generating a fluorescent Phen-DC3 derivative that retained the *in vitro* activity and cellular functions of the original compound and enabled live-cell fluorescence studies. This allowed us to unveil that the reason behind the surprisingly low effects of Phen-DC3 on cell viability is its inability to enter the nucleus. Instead, Phen-DC3 partially localizes in the mitochondria where it is affecting the mtDNA copy number. The developed synthetic platform was then used to link a cell-penetrating peptide to Phen-DC3. The penetrating peptide conjugation did not affect G4 selectivity and stabilization *in vitro* of the derivative compound but strongly increased the cellular effects. This is in line with an increased cellular and nuclear localization and opens up a whole new avenue for cell studies of G4 structures, *e.g*. by tuning the cell-penetrating peptide for use as delivery vehicles to also target specific tissues or tumors. Taken together, this shows that the developed synthetic platform is highly flexible and allows for the conjugation of both large and charged functionalities without affecting the ability to very selectively and strongly stabilize G4 DNA. The presented work thus disentangles important insights regarding the cellular localization and function of Phen-DC3 and describes a platform for various customized chemical biology studies of G4 DNA both *in vitro* and *in vivo*.

## Conflicts of interest

There are no conflicts to declare.

## Acknowledgements

Work in the Chorell lab was supported by the Kempe foundations (SMK-1632) and the Swedish Research Council (VR-NT 2017-05235). Work in the Wanrooij lab was supported by the Knut and Alice Wallenberg Foundation and the Swedish Research Council (VR-MH 2018-0278). MD was supported by HORIZON 2020-MSC individual fellowship (No 751474). We acknowledge Irene Martinez Carrasco and the Biochemical Imaging Centre at Umeå University and the National Microscopy infrastructure, NMI (VR-RFI 2019-00217), for providing assistance.

## Notes

### Competing Interest Statement

The authors have declared no competing interest.

